# D-glucuronyl C5-epimerase binds to EGFR to suppress kidney fibrosis

**DOI:** 10.1101/2024.11.25.625114

**Authors:** Xiaoqi Jing, Jun Wu, Jingru Ning, Xiaoyu Ding, Zhenyun Du, Xiaojiang Wang, Lulin Huang, Ran Wang, Changlin Mei, Kan Ding

**Affiliations:** Carbohydrate-Based Drug Research Center, CAS Key Laboratory of Receptor Research, State Key Laboratory of Drug Research, Shanghai Institute of Materia Medica, Chinese Academy of Sciences; Shanghai 201203 (China); Department of Nephrology, Changzheng Hospital, Second Military Medical University; Shanghai, 201203 (China); School of Chinese Materia Medica, Nanjing University of Chinese Medicine; Nanjing, 210023 (China); Drug Discovery and Design Center, State Key Laboratory of Drug Research, Shanghai Institute of Materia Medica, Chinese Academy of Sciences; Shanghai 201203 (China); Department of Natural Medicine, School of Pharmacy, Fudan University; Shanghai 201203 (China); Zhongshan Institute for Drug Discovery, Shanghai Institute of Materia Medica, Chinese Academy of Science; Zhongshan 528400 (China)

**Author notes:** Corresponding authors: Glycochemistry and Glycobiology Lab, Shanghai Institute of Materia Medica, Chinese Academy of Sciences; 555 Zu Chong Zhi Road, Pudong, Shanghai 201203, P. R. of China Kan Ding; Changlin Mei. These authors contribute to this work equally (Xiaoqi Jing, Jun Wu, Jingru Ning).

## Abstract

Renal tubular cells may actively participate in fibrosis processes leading to end-stage renal failure. However, which molecule play pivotal role in the fibrogenesis is still vague. Glucuronyl C5-epimerase (*Hsepi*, gene name, *Glce*) is a key enzyme that catalyzes biosynthesis of Heparan sulfate (HS) chains attached to HS proteoglycan which are ubiquitously located on cell membrane. Homozygous *Glce*-/- mice may cause embryonic lethality and multi-organ defects. However, whether *Glce* plays a key role in kidney fibrosis is unknown. Here, we show that *Glce* expression is significant attenuated in kidneys of patients with renal fibrosis and the animal models. Further study shows that renal tubular-specific *Glce* deletion in mice exacerbate kidney fibrosis while AAV-mediated overexpressing of *Glce* in UUO-treated mice may ameliorate kidney fibrosis associated with epithelial-mesenchymal transition (EMT) progress via the TGF-β/Smad2/3 signaling pathway. Mechanism study demonstrates that *Glce* protein may bind to EGFR to inactivate EGFR/ERK signaling and further impede TGF-β/Smad-driven EMT and renal fibrosis in Glce-/- and the wild type mice. Interestingly, the anti-fibrosis function is independent of *Glce* enzymatic activation.

These data uncover a novel function for *Glce* which plays a key role in kidney tissues against fibrosis.

## MAIN TEXT INTRODUCTION

Chronic Kidney Disease (CKD) affects one in ten people and will be the fifth highest cause of years of life lost worldwide by 2040 (*1*). Renal fibrosis, especially tubulointerstitial fibrosis (TIF), is a common feature of almost all progressive CKD, leading to a major determinant of renal insufficiency without treatment (*2*). During the development of chronic kidney disease, the accumulation of extracellular matrix (ECM) provides a springboard for progression and is regarded as a reliable predictor of prognosis. Despite the importance of this process, relatively little is known about its mechanism.

Recent studies have built a consensus on several key issues such as cell cycle arrest, defective cellular metabolism, and epithelial-to-mesenchymal transition (EMT) involved in the development and progression of kidney fibrosis (*3*). As one of the most critical roles, renal tubular cells have been demonstrated that they could actively participate in fibrosis processes. Based on this, studies suggested that targeting some aberrant expression of specific genes such as snail1, DsbA-L, and Tfam in tubular cells might directly or indirectly therapeutically ameliorate the kidney dysfunction(*4–6*). The research provided a novel insight for us and we believe that performing attention to renal tubular cells itself may bring advances to further clinical application in CKD treatment.

Heparan sulfate (HS) are linear glycan chains on HS proteoglycans (HSPGs) that are ubiquitously expressed on cell membrane and in the extracellular matrix of all tissues (*7*). HS chains not only are required for virus such as SARS-Cov-2 infection (*8, 9*), but also work as a co-receptor for fibroblast growth factor thought to engage pathogenesis of a spectrum of metabolic diseases, including obesity, non-alcoholic steatohepatitis, primary biliary cirrhosis, and CKD, etc(*7*). Evidence shows that HS plays a pivotal role in the fibrosis linked to chronic allograft dysfunction through binding growth factors (*10*). HS degradation enzyme, heparanase (*11*) or modification enzyme glucosaminyl-6-O-sulfotransferases functions in chronic renal fibrosis (*12*).

Glucuronyl C5-epimerase (*Hsepi*, gene name, *Glce*) is one of the modification enzymes involved in the biosynthesis of heparin and HS. A previous study found that the *Glce* could catalyze the C5-epimerization of the HS component, D-glucuronic acid (GlcA), into L-iduronic acid (IdoA), which provides internal flexibility to the polymer and forges protein-binding sites to ensure polymer function (*13*). As a highly conserved enzyme, *Glce* has been identified to exist widely not only between species but also in various types of organs, including the brain, lungs, kidneys, and many others (*14*). EEarlier studies indicated that targeting interruption of *Glce* in mice resulted in neonatal lethality accompanied by kidney agenesis, premature lung, and skeletal malformations, demonstrating that the single gene coded enzyme is essential for animal development (*15, 16*). Recently, we found that hepatic *Glce* deficiency led to impaired thermogenesis in adipose tissue and exacerbated high-fat diet (HFD)-induced obesity (*17*).

Despite its implication in the growth and development of various organs in animals, the exact physiological function of *Glce* remains largely unknow. Combined with previous studies suggesting that all *Glce*^-/-^ mice lacked kidneys and showed no overt abnormalities in other abdominal organs, while heparan sulfate structure alteration by its biosynthesis or degradation enzymes has impact on kidney chronic renal fibrosis (*11, 12*), we hypothesized that *Glce* might also have a role in renal fibrosis. In this report we examined the contribution of *Glce* to kidney development and its role in the pathogenesis of renal fibrosis. Using renal tubular epithelial cells and conditional knockout mice with clinical data from human renal tissues, we first show that *Glce* deficiency enhances epithelial-to-mesenchymal transition (EMT) in renal tubular epithelial cells *in vitro*, and genetic ablation of epithelial *Glce* results in aggravated fibrosis via EGFR/ERK pathway in the fibrosis models induced by unilateral ureteral obstruction (UUO). In addition, we demonstrate that renal-specific AAV-mediated *Glce* overexpression improves kidney fibrosis obviously and this process doesn’t rely on its catalytic isomerase activation. These findings reveal a new mechanism underlying the regulation of *Glce* activity during renal fibrosis and suggest the significance of preserving the basal *Glce* levels in the kidney as a therapeutic strategy to attenuate the progression of kidney fibrosis.

## RESULTS

### Glce was significantly downregulated in human and mouse fibrotic kidneys

To investigate the potential involvement of *Glce* in the pathogenesis of kidney fibrosis, we first examined *Glce* in renal biopsies from IgA nephropathy (IgA), lupus nephritis (LN), membranous nephropathy (MN), and diabetic nephropathy (DN) subjects compared with minimal-change nephrotic syndrome (MCNS) by immunohistochemistry (IHC) staining (Fig. 1A). Notably, the *Glce* positive rate was markedly reduced in renal biopsies from patients with different forms of chronic kidney disease compared with MCNS (Fig. 1B). Additionally, patients with advanced CKD showed a more accentuated decline of *Glce* positive rate except for the LN group (fig. S1A-H). We noticed that MCNS patients’ kidneys have far less interstitial fibrosis than other types of chronic kidney disease. (Fig. 1 A and C). To confirm the severity of kidney injury, we detected serum creatinine (Scr) and blood urea nitrogen (BUN) as the markers of renal functions (*18*). Results showed that the level of Scr and BUN in the MCNS groups was significantly lower than in others (Fig. 1D and E). Furthermore, we found the *Glce* positive rate was negatively correlated with the interstitial fibrosis area (Fig. 1F), serum creatinine (Fig. 1G), and blood urea nitrogen in all subjects (Fig. 1H). Considering that the *Glce* decline was significantly associated with nearly all prevalent CKD, we performed an additional analysis in mice subjected to unilateral ureteral occlusion (UUO) surgery or Folic acid (FA) treatment. In mouse kidney tissue, the results revealed that the mRNA and protein levels of *Glce* were reduced in the kidney of UUO and FA-treated mice over time (Fig. 1, J-M and fig. S2B and C). Furthermore, immunofluorescent results showed a significant reduction of *Glce* expression accompanied by an increase in α-SMA (*19*) in the kidneys of UUO mice and FA-treated mice (Fig. 1I).

**Fig. 1.**
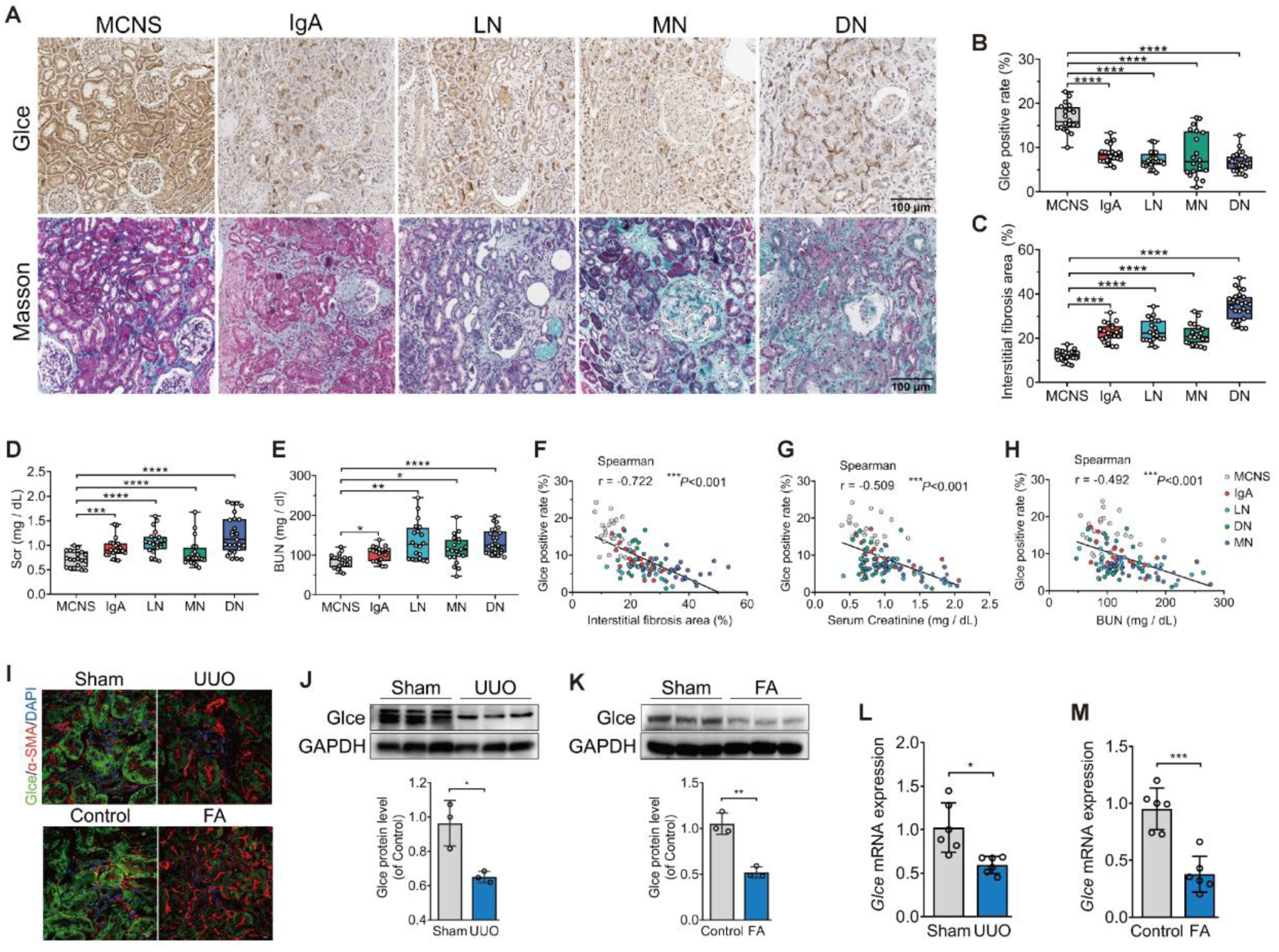
*Glce* was significantly downregulated in human and mouse fibrotic kidneys. **(A)** Representative photomicrographs of *Glce* protein using immunohistochemical staining (IHC) and Masson staining (MTS) in human renal tissues from minimal-change nephrotic syndrome (MCNS, n=20), IgA nephropathy (IgA, n=20), lupus nephritis (LN, n=20), membranous nephropathy (MN, n=20) and diabetic nephropathy (DN, n=25). Scale bars: 100 μm. (**B**) Positive rate of *Glce* protein with IHC staining and (**C**) interstitial fibrosis area in the renal biopsies from patients with different forms of chronic kidney disease. (**D**) Serum creatinine and (**E**) blood urea nitrogen (BUN) from patients with different forms of chronic kidney disease. (**F-H**) Negative correlation between *Glce* protein positive rate with IHC staining and interstitial fibrosis area (Spearman r = - 0.722, *p* < 0.001), Serum creatinine (Spearman r = - 0.509, *p* < 0.001), blood urea nitrogen (BUN) (Spearman r = - 0.492, *p* < 0.001) in all subjects, respectively. (**I**) Immunofluorescent staining of *Glce* (green), α-SMA (red) in kidneys from UUO- operated and FA-treated mice. Scale bars: 20 μm. (**J**) Immunoblots showed protein expression levels of *Glce* in the whole-kidney extract from WT mice 14 days after UUO (n = 6). (**K**) Immunoblots showed protein expression levels of *Glce* in the whole-kidney extract from WT mice a month after a single intraperitoneal injection of FA (250 mg/kg body weight) (n = 6). (**L**, **M**) *Glce* mRNA expression in the whole-kidney extract from WT mice following UUO or treated with FA (n = 6). GAPDH was used as loading control. Data are presented as the mean ±SEM. \**P* < 0.05; \*\**P* < 0.01; \*\*\**P* < 0.001; \*\*\*\**P* < 0.0001 by unpaired, 2-tailed Student’s *t* test (**J**, **K**, **L** and **M**), 1-way ANOVA with Dunnett’s *post hoc* tests (**B**, **C**, and **E**), Spearman’s correlation test (**F**, **G**, and **H**), and Kruskal-Wallis test (**D**).

### Glce deficiency aggravates renal failure and renal-specific AAV-mediated Glce overexpression improves kidney fibrosis

To determine the expression and distribution of *Glce* in the kidneys, kidney samples from patients and mice were stained. The results showed that *Glce* was predominantly present in tubular cells with a modest number of interstitial as well as glomerular cells. Interestingly, in both mouse and human tissues, most of the *Glce* protein was expressed in kidney tubular cells rather than in glomerular cells (fig. S2A). To elucidate the role of *Glce* in kidney fibrosis and injury, we deleted *Glce* in kidney tubules identified by tail genotyping (fig. S2D). Knockdown efficiency was verified by Western blotting (Fig. 2E) and immunofluorescence staining. Compared to Cdh16/*Glce*^flox/flox^ (*Glce*^-/-^) mice, control (Wild type, WT/*Glce*^flox/flox^) littermates mice showed no differences in α-SMA protein expression (fig. S2E and F) and maintained a higher kidney weight-to-body weight ratio at 8 weeks of age. This demonstrated the lack of *Glce* in the tubular cells led to significant retardation of kidney growth (fig. S2G). In addition, we interestingly found that following UUO surgery, some *Glce*^-/-^ mice displayed abnormal morphology in the contralateral kidney (Fig. 2A). We then measured serum creatinine and blood urea nitrogen to evaluate renal dysfunction. Compared to the control, the *Glce*^-/-^ mice group displayed significantly higher levels of Scr and BUN, reflecting that *Glce* knockdown induced more severe damage (Fig. 2B). Histological analysis indicated significant epithelial atrophy, dilated tubules, and interstitial fibrosis in the UUO group. *Glce* knockout mice markedly exacerbated UUO-induced tubular dilation, atrophy, and the notable accumulation of ECM (Fig. 2C). The ablation of *Glce* in tubular cells caused higher level of fibrosis markers in mRNA and protein expression compared to that in WT mice after UUO (Fig. 2D-F). Next, we overexpressed *Glce* in Cdh16/*Glce*^flox/flox^ mouse kidney *in vivo* using adeno-associated viral (AAV) vectors, after which the mice have subjected to UUO or Sham treatment after 6 weeks and sacrificed at 14 days post-UUO (fig. S2H). We first verify *Glce* protein was successfully expressed in the mice kidney by immunofluorescent analysis (fig. S2I). Interestingly, *Glce^-/-^*mice are protected against renal fibrosis as evidenced by an improved kidney morphology and a decrease in serum creatinine and BUN levels with UUO surgery after the AAV injection. (Fig. 2G and H). In addition, the histological staining revealed that the AAV-m*Glce* group had a more normal appearance with fewer deposition of the collagen fibers in kidney tissue of mice (Fig 2I). Further investigation manifested that overexpression of *Glce* prevents the higher fibrosis markers compared to the empty vector group mice (AAV-ZsG1) after UUO treatment (Fig. 2J-L).

**Fig. 2.**
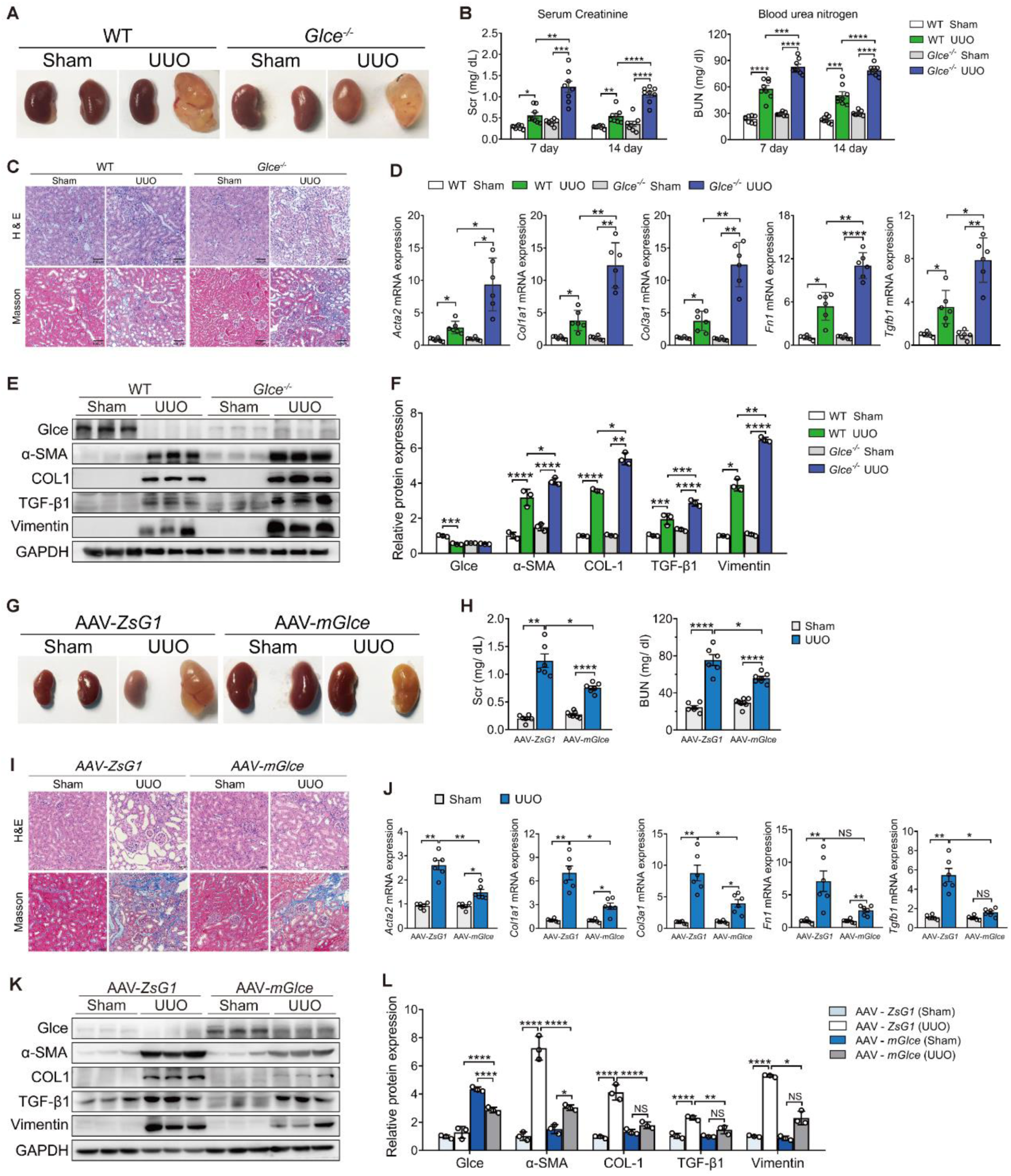
*Glce* deficiency aggravates renal failure and renal-specific AAV-mediated *Glce* overexpression improves kidney fibrosis. **(A)** Representative images of WT and *Glce*^-/-^ mice kidneys after sham operation or UUO surgery taken at 12 weeks of age. (**B**) Serum creatinine (Scr) and blood urea nitrogen (BUN) of WT and *Glce*^-/-^ mice at 7 and 14 days after sham operation or UUO surgery (n = 8). (**C**) Representative hematoxylin and eosin (H&E) staining, Masson staining of kidney sections from WT and *Glce*^-/-^ mice 14 days after sham operation or UUO surgery. Scale bar: 100 μm. (**D**) mRNA and (**E**, **F**) protein expression levels of fibrosis marker in WT and *Glce*^-/-^ mice at 14 days after sham operation or UUO (n = 3-6). (**G**) Morphological changes of kidneys in *Glce*^-/-^ mice with the sham operation or UUO surgery after AAV encoding vector plasmid (AAV-ZsG1) or full-length mouse *Glce* (AAV-m*Glce*) injection. (**H**) Scr and BUN levels of AAV-ZsG1 and AAV-m*Glce* mice after sham operation or UUO surgery (n = 6-7) (**I**) Representative hematoxylin and eosin (H&E) staining, Masson’s trichrome staining of *Glce*^-/-^ mice kidneys with sham operation or UUO surgery after the AAV injection. Scale bar: 100 μm. (**J**) mRNA and (**K**, **L**) protein expression levels of fibrosis marker in AAV-ZsG1 and AAV-m*Glce* mice following sham or UUO (n = 6). GAPDH was used as loading control. Data are presented as the mean ± SEM. \**P* < 0.05; \*\**P* < 0.01; \*\*\**P* < 0.001; \*\*\*\**P* < 0.0001 by 1-way ANOVA with Tukey or Dunnett’s *post hoc* tests (**B**, **D**, **F**, **H**, **J** and **L**).

### Tubule-specific *Glce* deletion exacerbates EMT progress via the TGF-β signaling pathway

To determine the role of *Glce* in the resistance to renal fibrosis, short hairpin RNA of *Glce* (sh*Glce* group) was employed to disrupt the *Glce* expression, and the control vector was used as negative control (NC group) in HK-2 cells. Intriguingly, a part of HK-2 cells from a cobblestone-like appearance to an elongated fibroblast-like shape by the knock-down of *Glce* expression, while overexpression of *Glce* had no effect on HK-2 cells (Fig. 3A). Additionally, we also observed the reduced mRNA and protein levels of *Glce* with increasing induction time by TGF-β treatment (Fig. S3A and B). This phenomenon suggests that *Glce* may affect the progression of Epithelial-mesenchymal transition (EMT) in renal tubular epithelial cells (*20–22*). The Western blotting analysis confirmed that the knockdown of *Glce* in HK-2 cells had profoundly increased expression of EMT marker genes at the protein level, including N-cadherin, Snail, and Slug as well as a diminution of E-cadherin (*23*) (Fig. 3B and C). In addition, the upregulation of EMT markers (*Snail1*, *Snail2*, and *Cdh2*) was significantly increased at the mRNA level in the *Glce^-/-^* mice kidneys compared to the WT group mice, while there is no difference of *Cdh1* between WT and *Glce^-/-^* mice after UUO (Fig. 3D). Immunoblots of whole-kidney tissue lysates also showed increased protein levels of EMT marker in *Glce^-/-^*mice compared with WT mice following UUO. Dedifferentiation markers E-cadherin were both decreased in the kidney tissues from WT and *Glce^-/-^*mice following UUO (Fig. 3E and F). To further confirm that *Glce* participates in the renal tubular EMT process, next we overexpressed *Glce* in HK-2 cells. The results showed that overexpression of *Glce* significantly attenuated TGF-β1- induced protein upregulation of Snail, Slug, and N-cadherin in HK-2 cells (Fig. 3G and H). In addition, the results showed that overexpression of *Glce* resulted in a marked downregulation of *Snail1*, *Snail2*, and *Cdh2*, as well as increased *Cdh1* compared to the control group following UUO (Fig. 3I). Consistent with the aforementioned results, immunoblots showed marked decreases of Snail, Slug, and N-cadherin in AAV-m*Glce* group following UUO (Fig. 3J and K). Collectively, these results demonstrate that *Glce* is critical to protect against EMT in response to kidney injury.

**Fig. 3.**
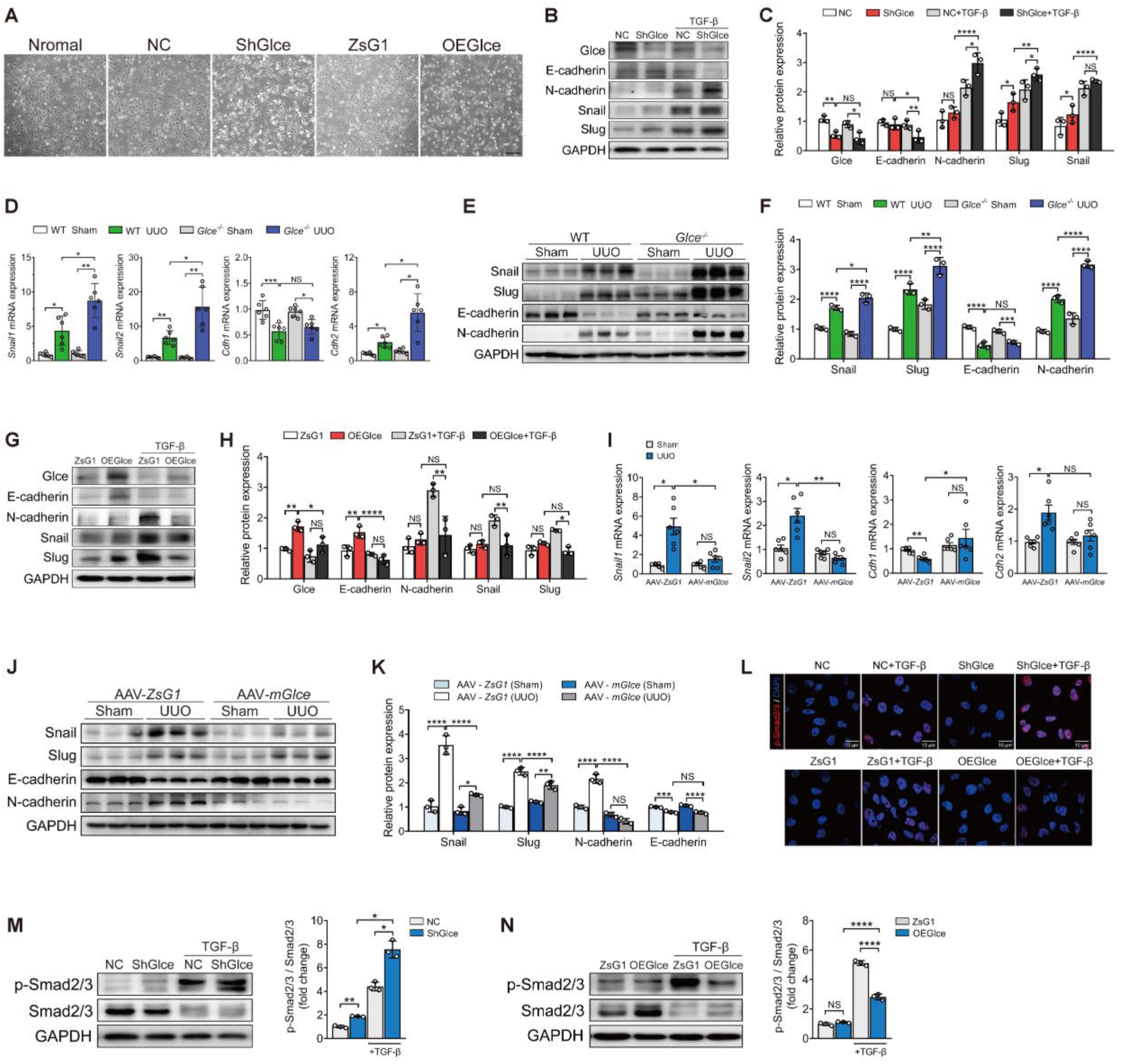
Tubule-specific *Glce* deletion exacerbates EMT progress via TGF-β signaling pathway. **(A)** Morphological changes in HK-2 cells upon incubated in serum-free medium (Normal) or stably transfected with empty vector (NC), *Glce* shRNA (Sh*Glce*), overexpression vector (ZsG1), and overexpression of *Glce* (OE*Glce*). Scale bar: 100 μm. (**B**) HK-2 cells were incubated separately with either serum-free medium or 10 ng/ml TGF-β1. Representative Western blots show levels of *Glce*, E- cadherin, N-cadherin, Slug, and Snail in HK-2 cells and the (**C**) corresponding quantification of them (n = 3). (**D**) Relative mRNA expression of *Snail1, Snail2, Cdh1, Cdh2* in WT and *Glce*^-/-^ mice after sham operation or UUO surgery (n = 6). (**E**) Protein expression levels of EMT marker in WT and *Glce*^-/-^ mice after sham operation or UUO and the (**F**) corresponding quantification of them (n = 3). (**G**) HK-2 cells were incubated with or without 10 ng/ml TGF-β1. Representative Western blots show levels of *Glce*, E-cadherin, N-cadherin, Slug, and Snail in HK- 2 cells and the (**H**) corresponding quantification of them (n = 3). (**I**) mRNA and (**J**, **K**) protein expression levels of EMT marker in AAV-ZsG1 and AAV-*mGlce* mice following sham or UUO (n = 6). (**L**) Immunofluorescence analysis for p- Smad2/3 (red) in HK-2 cells with or without 10 ng/ml TGF-β1. Scale bar: 10 µm. (**M**, **N**) Representative Western blots show levels of p-Smad2/3/Smad2/3 in HK- 2 cells and the corresponding quantification of them (n = 3). Data are presented as the mean ±SEM. \**P* < 0.05; \*\**P* < 0.01; \*\*\**P* < 0.001; \*\*\*\**P* < 0.0001 by 1-way ANOVA with Tukey or Dunnett’s *post hoc* tests (**C**, **D**, **F**, **H**, **I**, **K**, **M** and **N**).

According to the aforementioned, we speculated that *Glce* could mediate its differentiation effects through regulating signaling effectors downstream of TGF-β1.

We then examined the activities of the TGF-β-Smad2/3 pathway, which are known to have a critical role in the EMT of renal tubular cells and the progression of kidney fibrosis (*24, 25*). Western blotting analysis showed that tubule-specific *Glce* deletion promotes the phosphorylation of Smad2/3 and can be reversed by overexpression of *Glce* (Fig. 3M and N). The same result also obtained by the immunofluorescence image (Fig. 3L). Because TGF-β1-mediated Smad2/3 activation is strictly dependent on transmembrane TGF-β receptor type I (TGFBR1). We then designed an experimental procedure and treated the animals with a TGFBR1 inhibitor (SB431542) (Fig. S3C). Remarkably, in SD mice treated with SB431542, the kidney function indices (Scr, BUN) improved after UUO, whereas no discernible change was seen in WT mice. (Fig. S3D). Additionally, areas of collagen deposition and tubular necrosis became smaller after treatment of SB431542 in the *Glce*^-/-^ mice group with UUO surgery, while the WT mice did not show signs of improvement compared with *Glce*^-/-^ mice and some individual mice even deteriorated (Fig. S3E). Analogously, immunohistochemistry results showed that protein expression of α-SMA, and COL1 decreased in the kidneys of *Glce*^-/-^ mice after UUO, while no significant changes were found in WT mice (fig. S3F). Western blot indicated that SB431542 treatment suppressed the phosphorylation of Smad2/3, and the protein level of N-cadherin, Snail, and Slug in both groups of mice (fig. S3H and S3I). As the disease progressed, obstructed kidneys of the WT and *Glce*^-/-^ mice maintained the lower *Cdh1* expression levels and higher expression levels of *Snail1*, *Snail2,* and *Cdh2* genes compared with the control mice, and treatment with SB431542 could lead to the reverse trend in *Glce*^-/-^ mice but have no effect on *Snail2* and *Cdh1* in WT mice. (Fig. S3G).

### The Glce binds to EGFR to suppress its activation and impact on renal fibrosis

The previous sections have shown that *Glce* played vital role in the regulation of EMT via the TGFβ1/Smad pathway. However, upon further evaluation, we found there’s no direct interaction between *Glce* and TGFBR1 (Unpublished data). It has been reported that the enhanced activation of the EGF receptor (EGFR) associated with the development and progression of renal fibrosis and genetic or pharmacologic blockade of EGFR could inhibit renal fibrosis(*26*). To explore whether EGFR might be a target of the *Glce* protein, we firstly carried out the surface plasmon resonance (SPR) experiment, intriguingly, we eventually found that the *Glce* protein indeed had a strong binding interaction with the intracellular domain of EGFR *in vitro* however did not bind to the EGFR extracellular domain. As a positive control, N-Sulfated K5 polysaccharide, a classical substrate (*27*) for the *Glce* enzyme was used (Fig. 4A). Next, to further confirm the result and analyze the binding details, docking analysis was performed. As shown in Fig. 4B (Up panel), the cytoplasmic domain of EGFR is predicted to bind into a deep hydrophobic groove of *Glce* protein that is spanned by (α/α)4-barrel domain and the β-sandwich domain. The majority of the buried surface area between the docking *Glce*-EGFR complex is calculated as 2363 Å2. The structure indicated by the dashed lines serves as the predicted site that forming key interactions (Down panel). The *Glce* protein might form 11 hydrogen bonds with EGFR (black dashed lines), including N255-T969’, M402-T969’, T349-E980’, T349-R975’, N393-D984’, R266-D984’, W347-D982’, S265-N747’ and R266-M983’. Meanwhile, the *Glce* protein might also form salt bridges between 4 residue pairs, including D297-R975’, R266-D984’, K264- D746’, K299-D982’ (red dashed lines). The detailed distance information is provided in Supplemental Table 1. We next wonder if *Glce* colocalizes with EGFR in the renal tubular cells, as shown in Fig. 4C, *Glce* displayed a strong co-localization with EGFR in each group except the Sh*Glce* group in HK-2 cells. To confirm these interactions, we also performed co-immunoprecipitation (Co-IP) assays. Although the precipitate confirmed the binding of *Glce* to EGFR in HK-2 cells, however, the interaction between *Glce* and EGFR was markedly decreased when the EGFR was activated by EGF (Fig. 4D). Thus, we speculated that *Glce* might be implicated in the activation of EGFR. To investigate whether *Glce* regulates the phosphorylation of EGFR, we analyzed the major phosphorylation sites of EGFR at Tyr1045, Tyr1068, Tyr1148, and Tyr1173/Tyr1248 by immunoblot. Immunoblots demonstrated that *Glce* knockdown increased the phosphorylation of EGFR at Tyr1068 while having no measurable effect on other tyrosine phosphorylation sites (fig. S4A). Consistently, immunofluorescence (IF) staining showed the increased level of the phosphorylated EGFR in Sh*Glce* group rather than other groups (Fig. 4G). Previous studies have proved that activation of the EGFR/MAPK pathway led to the progression of renal fibrosis (*28*). To study the effect of *Glce* on the MAPK pathway, we assessed the activation of the pathway in the *Glce* knockdown cell lines. The results showed *Glce* knockdown did not significantly alter the expression of p-p38, and p-JNK, but markedly activate the phosphorylation of ERK1/2 and its downstream signaling pathways (Fig. 4E and F, and fig. S4B and C). We then overexpressed *Glce* and detected the expression of the MAPK pathway, but no significant change was observed compared to the vector control group (Fig. 4H and I). The results confirmed that the *Glce* knockdown might activate EGFR/ERK pathways, however, the overexpression of *Glce* did not affect EGFR phosphorylation in HK-2 cells. Immunoblots of whole-kidney tissue lysates also showed reduced protein levels of the p-EGFR(Tyr1068) and p-ERK1/2 in AAV-m*Glce* mice compared with AAV- ZsG1 mice following UUO (Fig. 4J and K). In addition, protein coimmunoprecipitation experiments in AAV-m*Glce* mice kidneys showed physical binding between *Glce* and EGFR, while the interaction between them was decreased in UUO-induced fibrotic kidneys (Fig. 4L).

**Fig. 4.**
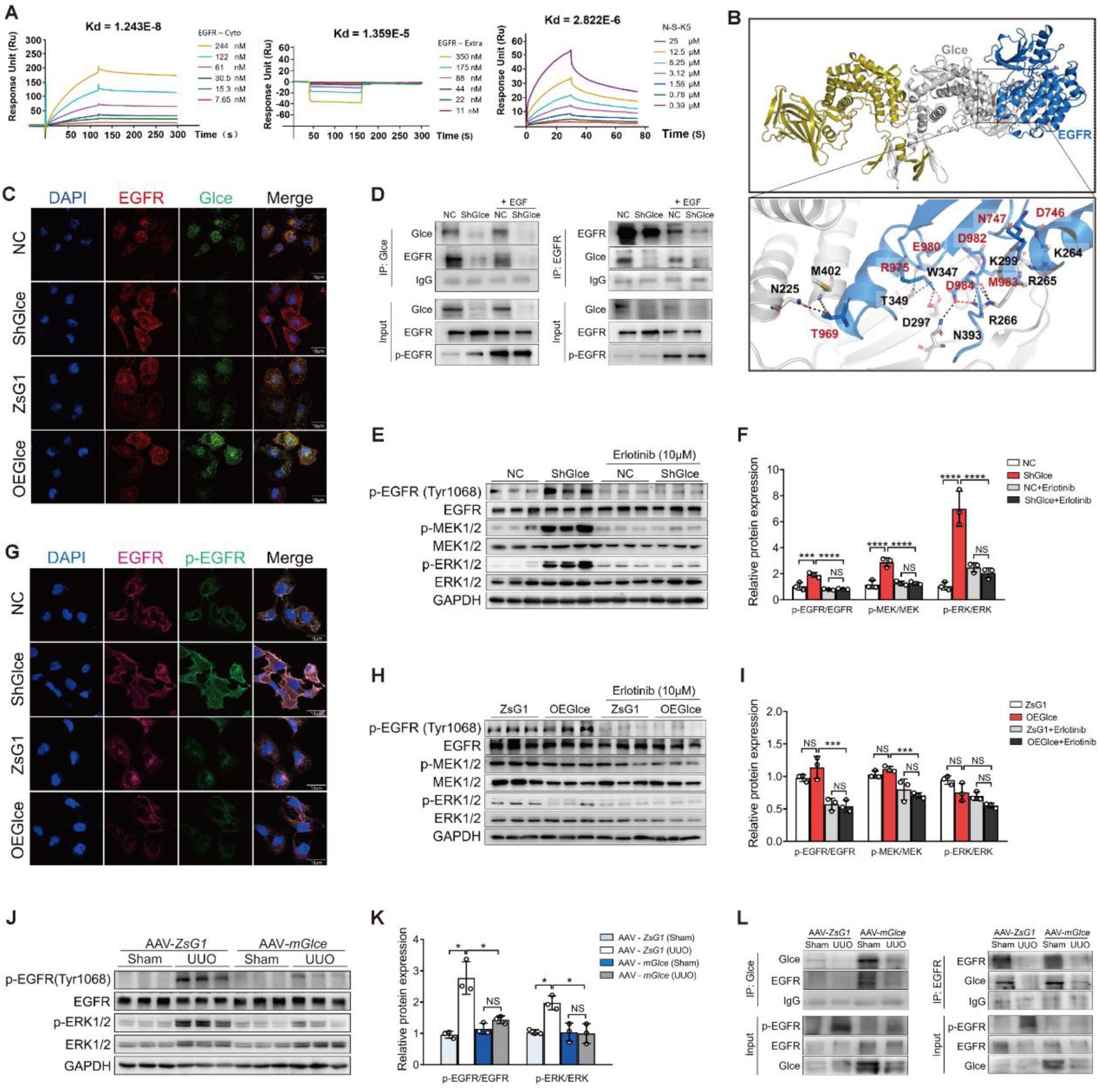
The *Glce* binds to EGFR to suppress its activation and impact on renal fibrosis. **(A)** SPR test showed direct binding between *Glce* and EGFR proteins. Sensorgrams of intracellular EGFR protein (7.65-244 nM) binding to *Glce* protein (left), extracellular EGFR protein (11-350 nM) binding to the *Glce* protein (middle), positive control polysaccharide N-S-K5 (0.39-25 μM) binding to *Glce* (right). (**B**) The docking model of the *Glce*-EGFR complex. The intracellular domain of EGFR was shown in a blue cartoon representation, respectively. Two monomers of h*Glce* were shown in gold and silver, respectively. The binding interface was marked with a black rectangular box and the close-up view of the interactions was shown in the down panel. The key residues involved in *Glce* (black) and EGFR (red) interactions were shown as sticks and colored silver and blue, respectively. Salt bridges and hydrogen bonds were shown as red and black dashed lines, respectively. (**C**) Immunofluorescence co-localization analysis for *Glce* (green), EGFR (red) in HK- 2 cells transfected with different kinds of plasmids. Scale bar: 10 µm. (**D**) Co-IP analysis of the interactions between *Glce* protein and EGFR in HK-2 cells. GAPDH was used as input loading control. Data are representative of at least two independent experiments. (**E**) Representative western blots of p-EGFR (Tyr1068), EGFR, p-MEK1/2, MEK1/2, p-ERK1/2, ERK1/2 in HK-2 cells transfected with sh*Glce* and vector control (NC) followed by Erlotinib (10 μM) and (**F**) protein quantification (n = 3). (**G**) Immunofluorescence co-localization analysis for p- EGFR (green), EGFR (rose) in HK-2 cells transfected with different kinds of plasmids. Scale bar: 10 µm. (**H**) Representative western blots of p-EGFR (Tyr1068), EGFR, p-MEK1/2, MEK1/2, p-ERK1/2, ERK1/2 in HK-2 cells transfected with *Glce*-overexpression plasmid and vector control (ZsG1) followed by Erlotinib (10 μM) and (**I**) protein quantification (n = 3). GAPDH was used as loading control. (**J**) Representative Western blots of p-EGFR (Tyr1068), EGFR, p-ERK1/2, ERK1/2 and (**K**) protein quantification in the kidneys of *Glce*^-/-^ mice with sham operation or UUO surgery after AAV encoding full-length *Glce* injection (n = 3). (**L**) Co-IP analysis of the interactions between *Glce* protein and EGFR in the kidneys of *Glce*^-/-^ mice with sham operation or UUO surgery after the AAV injection. GAPDH was used as input loading control. Data are representative of at least two independent experiments. Data are presented as the mean ±SEM. \**P* < 0.05; \*\**P* < 0.01; \*\*\**P* < 0.001; \*\*\*\**P* < 0.0001 by 1-way ANOVA with Tukey or Dunnett’s *post hoc* tests (**F**, **I** and **K**).

### Inhibition of EGFR Signaling pathway ameliorates TGF-β/Smad-driven EMT and renal fibrosis in the *Glce*^-/-^ and WT mice

Previous studies had shown that TGF-β-mediated tissue fibrosis relies on a persistent feed-forward mechanism of EGFR/ERK activation (*29*). To explore the effects of *Glce* on pathogenesis and progression of renal fibrosis *in vivo* through the EGFR/ERK signaling pathway, an EGFR inhibitor erlotinib was employed following UUO surgery (Fig. 5A). Histological analysis indicated significant epithelial atrophy, dilated tubules, and interstitial fibrosis after UUO, of which *Glce*^-/-^ mice performed more severe changes. After erlotinib administration for 14 days, the renal damage was significantly ameliorated (Fig. 5B). Additionally, the biochemical assays demonstrated that levels of serum creatinine and blood urea nitrogen was increased by varying degrees in UUO- treated animals, however the levels were significantly decreased in both groups that receiving erlotinib treatment (Fig. 5C). Western blot analysis from kidney lysates confirmed the protein levels of p-EGFR(Tyr1068) and p-ERK1/2 were increased in both WT and *Glce*^-/-^ mice following UUO. However, the extent of this was reduced with erlotinib treatment (Fig. 5D and E). As shown in Fig. 5F, the protein levels of COL1, α-SMA, and vimentin were increased following UUO, whereas erlotinib administration led to a marked reduction of them in both WT and *Glce^-/-^* mice. Compared with WT mice, UUO-treated *Glce*^-/-^ mice showed further increased protein expression. Consistently, enhanced expression of TGF-β1, p-Smad2/3, and its downstream EMT markers were markedly decreased after administration of erlotinib (Fig. 5G). Therefore, we concluded that the genetic deletion of *Glce* activated the EGFR signal pathway. Subsequently, aberrant EGFR activation resulted in increased expression of TGF-β1 and then exacerbated the EMT process via SMAD-dependent pathways in kidneys.

**Fig. 5.**
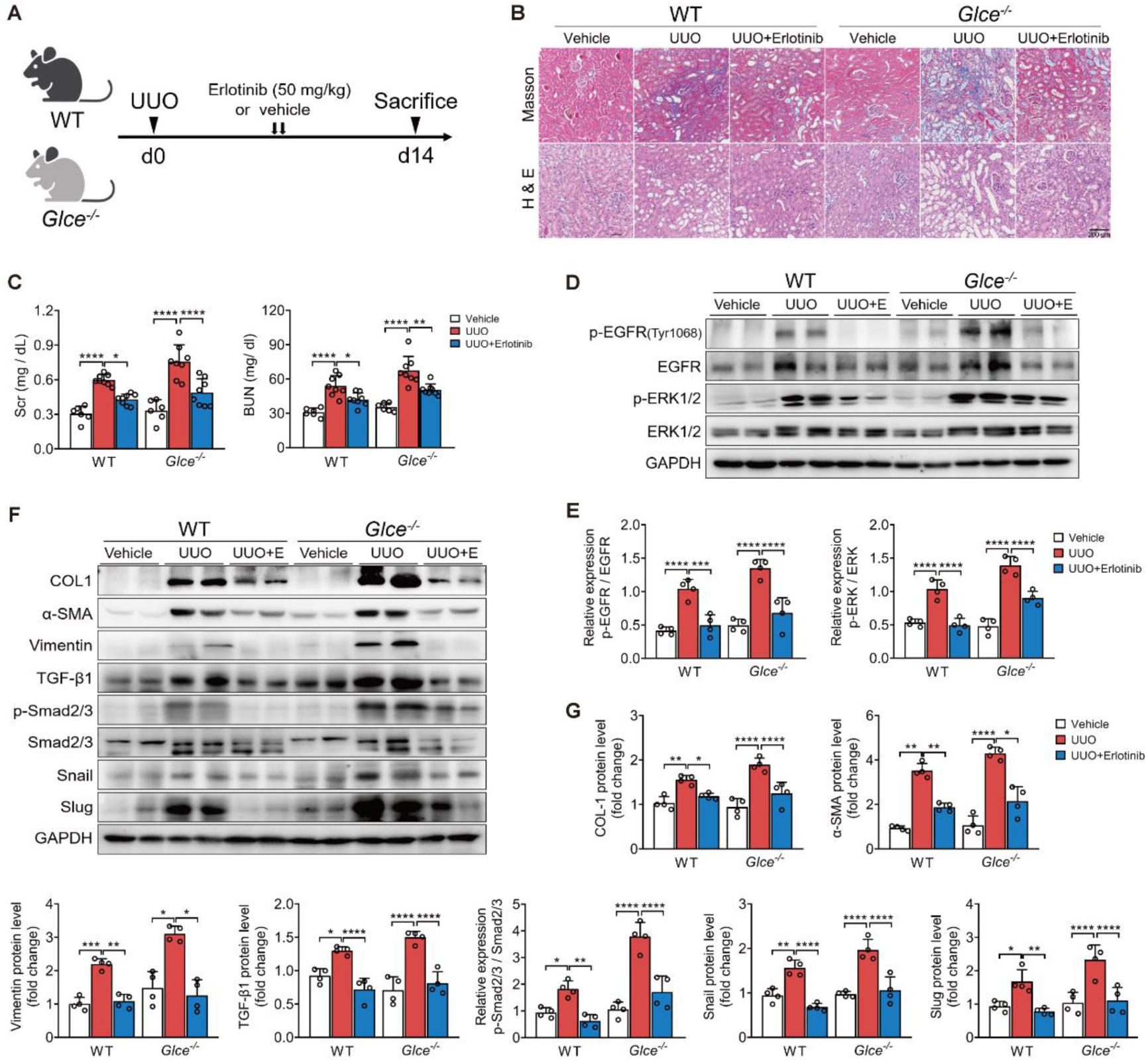
Inhibition of EGFR Signaling pathway ameliorates TGF-β/Smad-driven EMT and renal fibrosis in the *Glce^-/-^* and WT mice. **(A)** A schematic diagram of treatment with erlotinib in the UUO-induced mice. (**B**) Representative hematoxylin and eosin (H&E) staining, Masson staining of WT and *Glce*^-/-^ mice kidneys following erlotinib treatment 14 days after UUO. Scale bar: 100 μm. (**C**) Serum creatinine and blood urea nitrogen of WT and *Glce*^-/-^ mice following erlotinib treatment 14 days after UUO. Sham group (n = 6), UUO group (n = 8), UUO + Erlotinib group (n = 8). (**D, E**) Western blot analysis of p-EGFR (Thr1068), EGFR, p-ERK1/2, ERK1/2 and GAPDH in whole-kidney lysates from WT and *Glce*^-/-^ mice 14 days after UUO and the corresponding quantification of p- EGFR/EGFR and p-ERK1/2/ERK1/2 (n = 4). (**F, G**) Representative Western blots of Collagen 1, α-SMA, Vimentin, TGF-β1, p-Smad2/3, Smad2/3, Snail, Slug and protein quantification in WT and *Glce*^-/-^ mice kidneys following erlotinib treatment 14 days after UUO (n = 4). Data are representative of at least three independent experiments. Data are presented as the mean ±SEM. \**P* < 0.05; \*\**P* < 0.01; \*\*\**P* < 0.001; \*\*\*\**P* < 0.0001 by 1-way ANOVA with Tukey or Dunnett’s *post hoc* tests (**C**, **E** and **G**).

### Adeno-associated virus encoding mutant Glce orthotopically injected to Glce^-/-^ mice still can ameliorate renal fibrosis

According to SPR results, the *Glce* protein may robustly interact with EGFR’s intracellular region. Molecular docking analysis revealed that the potential EGFR binding regions in human *Glce* protein are mostly located in its classical β-sandwich domain and did not include the classical enzyme active sites of *Glce*. Therefore, to determine whether the enzymatic function of *Glce* was essential for EGFR activation and TGF-β1-induced EMT in the pathogenesis of renal fibrosis, we next chose three sites that were considered crucial for the enzyme activity including Y500, Y560, and Y578, respectively for mutational studies (*16*). HK-2 cells with stable *Glce* knockdown were transfected with plasmids containing mutant genes. The protein expression level of p-EGFR, p-MEK1/2, and p-ERK1/2 was markedly higher in the *Glce* knockdown HK-2 cells, while mutant *Glce* showed no effect on the EGFR/ERK pathway (Fig. 6A). In previous studies, it was suggested that HS is biosynthesized in the Golgi network, where the polysaccharide chain is polymerized and modified by a series of Golgi-located enzymes (*30*). Interestingly, we found that the mutant *Glce* was expressed successfully and distributed in the cytoplasm of renal tubular cells besides Golgi (Fig. 6B). To understand the therapeutic potential of mutant *Glce*, we constructed the adeno-associated virus vectors encoding GFP as well as all three mutant sites (AAV-mut*Glce*) simultaneously and then injected them orthotopically into the kidney of *Glce*^-/-^ mice. Empty vector plasmid (AAV-ZsG1) was utilized as a control. Immunofluorescence staining results showed that the mutant *Glce* was overexpressed successfully in renal tubular epithelial cells (Fig. 6C). Compared with those injected with empty vector plasmid (AAV-ZsG1), the overexpression of mutant *Glce* (AAV- mut*Glce*) resulted in a lower level of Scr and BUN following UUO (Fig. 6D). Overexpression of mutant *Glce* reduced the extent of renal fibrosis compared with AAV- ZsG1 mice (Fig. 6E). Immunoblots showed reduced fibrosis protein levels in AAV- mut*Glce* mice compared with AAV- ZsG1 mice following UUO (Fig. 6F and G). Additionally, expression of TGF-β1 and EMT markers were markedly lower in AAV- mut*Glce* mice compared with AAV- ZsG1 mice subjected to UUO (Fig. 6H and I). Similarly, Western blot analysis revealed that overexpression of mutant *Glce* induced a marked decrease in phosphorylation of EGFR/ERK signaling pathway compared with vector control mice in UUO-treated kidneys (Fig. 6K and L). Furthermore, the immunofluorescence analysis showed co-location of EGFR and mutant *Glce* in kidney tissues from AAV-mut*Glce* mice (Fig. 6J). Taken together, AAV- mut*Glce* mice challenged with UUO showed better tubular health and renal function, and less interstitial fibrosis.

**Fig. 6.**
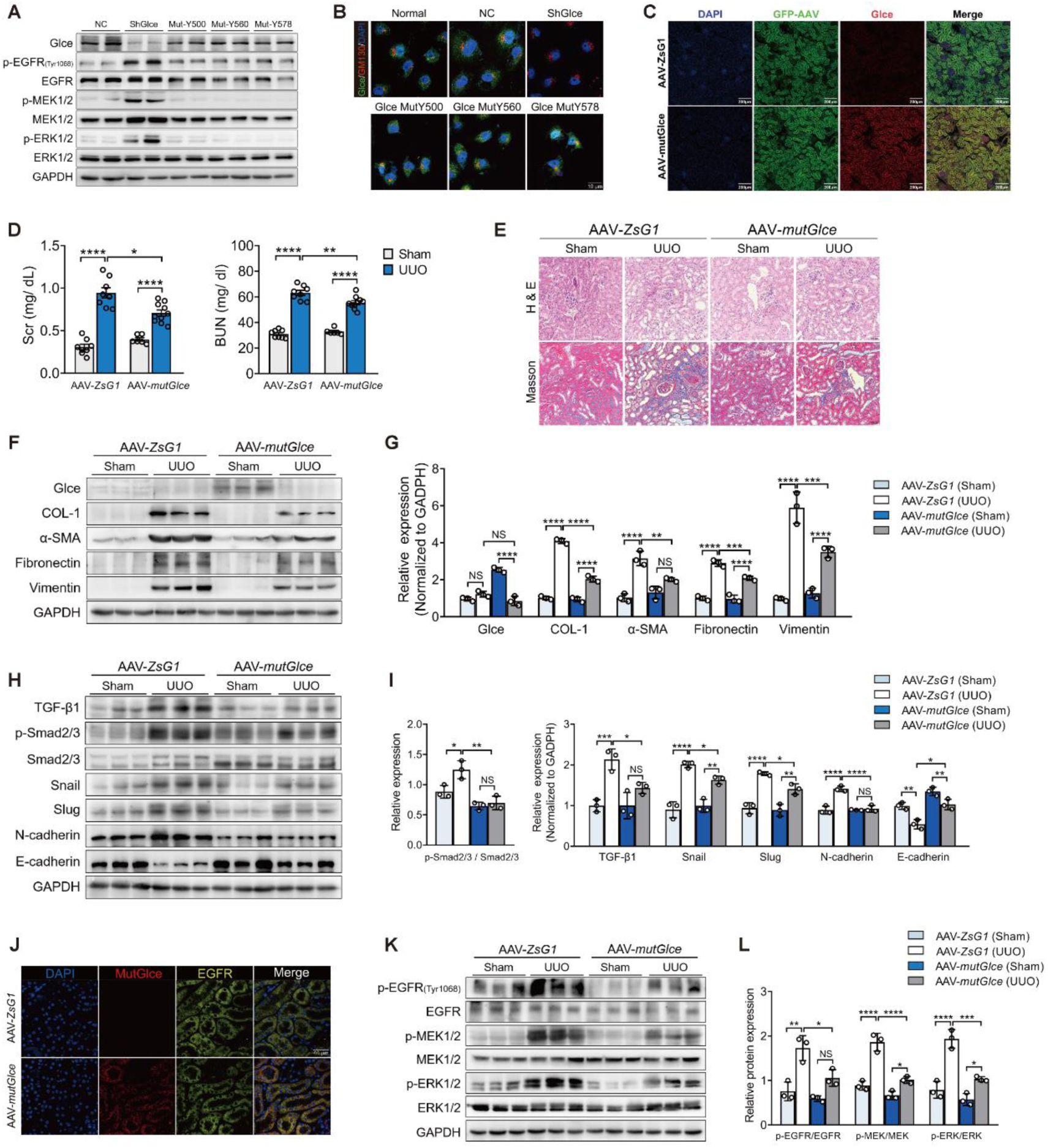
Adeno-associated virus encoding mutant *Glce* orthotopically injected to *Glce*^-/-^ mice still can ameliorate renal fibrosis. **(A)** Representative Western blots of *Glce*, p-EGFR (Thr1068), EGFR, p-MEK1/2, MEK1/2, p-ERK1/2, ERK1/2 in HK-2 cells transfected with plasmids encoding control vector (NC), sh*Glce* and mutant *Glce* (Y500F, Y560F, Y578F). GAPDH was used as loading control, data are representative of two independent experiments. (**B**) Immunofluorescent staining of *Glce* (green), GM130 (red) in HK- 2 cells with different genotypes. Scale bar: 10 μm. (**C**) Immunofluorescence staining of *Glce* (red) and GFP (green) in the mice 6 weeks after AAV encoding full-length mutant *Glce* injection. Scale bar: 200 μm. (**D**) Serum creatinine and blood urea nitrogen of *Glce*^-/-^ mice with sham operation or UUO surgery after AAV encoding full-length mutant *Glce* injection. AAV-ZsG1 Sham group (n = 8), UUO group (n = 8); AAV-m*Glce* Sham group (n = 6), UUO group (n = 10). (**E**) Representative hematoxylin and eosin (H&E) staining, Masson staining of *Glce^-/-^* mice kidneys with sham operation or UUO surgery 6 weeks after AAV-ZsG1 or AAV-mut*Glce* injection. Scale bar: 100 μm. (**F**, **G**) Representative Western blots of *Glce*, Collagen 1, α-SMA, Fibronectin, Vimentin and protein quantification in the kidneys of *Glce*^-/-^ mice with sham operation or UUO surgery after AAV-ZsG1 or AAV-mut*Glce* injection (n = 3). (**H**, **I**) Representative Western blots of TGF-β1, p-Smad2/3, Smad2/3, Snail, Slug, E-cadherin, N-cadherin and protein quantification in the kidneys of *Glce*^-/-^ mice with sham operation or UUO surgery after the AAV injection (n = 3). (**J**) Immunofluorescence co-localization analysis for mutant *Glce* (red), EGFR (yellow) in the kidneys of *Glce*^-/-^ mice after AAV- ZsG1 or AAV-mut*Glce* injection. Scale bar: 20 μm. (**K**, **L**) Representative Western blots of p-EGFR (Thr1068), EGFR, p-MEK1/2, MEK1/2, p-ERK1/2, ERK1/2 in the kidneys of *Glce*^-/-^ mice with sham operation or UUO surgery after AAV injection (n = 3). Data are representative of at least three independent experiments unless indicated otherwise. Data are presented as the mean ±SEM. \**P* < 0.05; \*\**P* < 0.01; \*\*\**P* < 0.001; \*\*\*\**P* < 0.0001 by 1-way ANOVA with Tukey or Dunnett’s *post hoc* tests (**D**, **G**, **I** and **L**).

## DISCUSSION

Kidney fibrosis is a characteristic feature of all forms of chronic kidney disease. As the fibrosis is a complex process involving different mechanisms and factors, so far, the therapy for renal fibrosis is quite limited in clinic (*31*). As is well known, *Glce* is a highly evolutionarily conserved enzyme expressed in various organs in the body (*32*). On this basis, we constructed renal tubular-specific *Glce* knockout mice and observed the phenotype during mouse development. Interestingly, these mice did not die embryonically and showed no obvious difference in activity and appearance compared with the control mice. However, the renal weight/body weight ratio measurements after sacrifice confirmed that renal growth retardation occurred in *Glce*^-/-^ mice. More seriously, the individual *Glce* gene-deficient mice even displayed unilateral absence of kidneys. Subsequently, we were surprised to find that *Glce* levels were lower in patients with various types of CKD, and its levels negatively correlated with the degree of fibrosis at different stages. This result suggested that the *Glce* might be related to renal fibrosis. We then examined the typical fibrosis signaling pathway including TGF- β/Smad, Wnt/β-catenin, Hedgehog, PI3K/Akt/mTOR and Notch in the *Glce* knock out mice (*33–37*). Whereas, the result indicated that except TGF-β/Smad knockout of *Glce* did not induce the change in these signaling pathways after *Glce* deletion compared to WT mice.

Now that we have shown that *Glce* is not the direct cause for fibrosis, instead, we observed that *Glce* knockdown promoted epithelial–mesenchymal transition in the HK-2 cells. TGF-β1 is considered a key mediator of EMT by inducing transcription of several mesenchymal genes and increasing the activity of EMT transcription factors via SMAD factors in renal epithelial cells (*20, 38*)^,^ Our study revealed that *Glce* deficiency in renal tubular cells promoted the activation of TGF-β/Smad pathways predominantly associated with TGFBR1 rather than Smad3 and thereby exacerbated EMT progress. However, inhibition of TGF-β/Smad pathways partially relieved the EMT process and renal fibrosis in *Glce*^-/-^ mice after UUO. Moreover, further investigation indicated that *Glce* bind to neither TGF-β1 nor TGFB1 receptor. Surprisingly, we found that *Glce* could bind strongly to EGFR. Our results indicated that overexpression of *Glce* prevents the abnormal activation of the EGFR pathway and improved renal tubulointerstitial fibrosis. EGFR is a transmembrane glycoprotein that is composed of an extracellular region, a transmembrane domain, and an intracellular tyrosine kinase domain (*39*). By using SPR, we observed that *Glce* binds with high affinity to the intracellular domain of EGFR. We consider that their binding may primarily influence the EGFR kinase domains activation, and indeed the docking prediction also confirmed it, which is consistent with the results observed in our *in vitro* and *in vivo* experiments. Thus, we believed that the *Glce*-EGFR binding effects act as an ‘on-off’ like switch to regulate the activation of EGFR in the kidneys.

More evidences had already suggested that heparan sulfate (HS) played pivotal role in various pathological fibrosis (*40*). Therefore, we speculated that the renoprotection of *Glce* relies on the function catalyzing the formation of HS. Surprisingly however, the results indicated that the *Glce* enzyme mutation in tubular cells not only did not result in the activation of the EGFR pathway but also still ameliorate renal fibrosis and the TGF-β-mediated EMT process in the UUO-induced *Glce*^-/-^ mice. Additionally, we observed that *Glce* also distributed near the cellular membrane and colocalized with EGFR besides Golgi. Although most enzymes modulate fibrosis by their catalytic mechanism, it has also been shown that the non-enzymatic function of proteins was implicated in the progression of fibrosis. Recently, we showed that *Glce* protein might bind to GDF15 to maintain energy metabolism balance in a non-enzymatic function (*17*). Based on this, we infer that in addition to playing an epimerase function in the Golgi apparatus, *Glce* can also regulate EGFR thereby activating distinct signaling pathways by a non-enzymatic mechanism. Overall, the protective effects of *Glce* against renal fibrosis suggested its multifunctional roles and we were delighted to discover a novel function of *Glce*.

In conclusion, this study for the first time, provide evidences for the important role of *Glce* played in the maintenance of kidney function and against renal fibrosis. We uncover that *Glce* can bind to the intracellular domains of EGFR in normal tubular cells. Tubule-specific *Glce* loss causes the activation of EGFR signaling as well as the TGF- β-dependent EMT process, leading to more severe tubulointerstitial injury and fibrosis after UUO. Particularly, one novel aspect in this work is the finding that the nephroprotective effect of *Glce* distinguishes itself from the currently well-known enzymatic function. Therefore, our data provide a novel potential target for the treatment of chronic kidney disease.

## MATERIALS AND METHODS

### Study Design

The objective of this study was to determine the protective role of renal D-Glucuronyl C5-Epimerase (Glce) in the pathogenesis of kidney fibrosis. Using examined immunohistochemistry (IHC) staining, we firstly identified the reduction of Glce in renal biopsies from different types of CKD patients compared with minimal-change nephrotic syndrome (MCNS), which was closely related to the degree of fibrosis deterioration. So, we hypothesized that Glce reasonably plays an important role in the progression of kidney fibrosis. To test it, we then chose UUO and FA-treated mice as the model of CKD. Combined with the Cdh16/Glceflox/flox (Glce-/-) mice model, we further evaluated the effect of Glce on the renal function. Next, by performing the RT-PCR and western blotting analysis, we showed that tubule-specific Glce deletion exacerbates Epithelial-mesenchymal transition (EMT) progress via the TGF-β signaling pathway in vitro experiments using HK-2 cells.

To explore the molecular mechanism of the promotion of EMT and renal fibrosis caused by Glce deletion, we performed the SPR and Co-IP assay which assessed the potential interaction of Glce with EGFR. Next, to further confirm the result and analyze the binding details, a molecular model of interaction sites for Glce protein binding to EGFR was performed by docking analysis. To further determine if Glce colocalizes with EGFR in the renal tubular cells, we performed the confocal immunofluorescence analysis. To study the effect of Glce on the MAPK pathway, we assessed the activation of the pathway in the Glce knockdown and overexpression cell lines. Additionally, to explore the effects of Glce on pathogenesis and progression of renal fibrosis in vivo through the EGFR/ERK signaling pathway, an EGFR inhibitor erlotinib was employed following UUO surgery. Moreover, to determine whether the enzymatic function of Glce was essential for EGFR activation and TGF-β1-induced EMT in the pathogenesis of renal fibrosis, we next chose three sites that were considered crucial for the enzyme activity including Y500, Y560, and Y578, respectively for mutational studies. To understand the therapeutic potential of mutant Glce, we constructed the adeno-associated virus vectors encoding GFP as well as all three mutant sites (AAV-mutGlce) simultaneously and then injected them orthotopically into the kidney of Glce-/- mice. Empty vector plasmid (AAV-ZsG1) was utilized as a control. Within the littermate groups, animals were selected at random for each experimental group. The number of biological replicates is indicated in the figure legends.

### Human renal biopsy samples

Patients were recruited through the Department of Nephrology in Shanghai Changzheng Hospital in Shanghai. All clinical records involved in this study obtained Institutional Review Board approval and the raw data used in the present study were obtained from clinical records of participating patients and were included in a database after anonymization (2022SL064). The patients were grouped into minimal-change nephrotic syndrome (MCNS), IgA nephropathy (IgA), lupus nephritis (LN), membranous nephropathy (MN), and diabetic nephropathy (DN) according to different features of diseases. Several trained nephrologists evaluated clinical imaging to define the specific pathological types of renal disease. Clinical patient blood biochemical parameters were extracted from the custom system of Shanghai Changzheng Hospital.

### Mouse models

All animals were maintained in the core animal facility and approved by the Institution Animal Care and Use Committee at Shanghai Institute of Materia Medica. Eight- to ten-week-old C57BL/6 mice were ordered from Shanghai SLAC Laboratory Animal Company (Shanghai, China). All experiments were conducted on group-housed (4–5 animals per cage) animals under a 12-hour light/dark cycle with food and water freely accessible. Animal studies were approved by the *IACUC* of Shanghai Institute of Materia Medica (*IACUC* no. 2019-06-DK-79; 2019-06-DK-80).

The floxed *Glce* mice and Cdh16-cre transgenic mice were generated and purchased from the Model Animal Research Center of Nanjing University (Nanjing, China). Floxed *Glce* mice were backcrossed with C57BL/6 mice for more than 8 generations to produce congenic strains. Then C57BL/6 *Glce* ^flox/flox^ mice were crossed with mice expressing Cre recombinase (Cre) to generate renal tubular-specific *Glce* knockout mice (*Glce*^-/-^ mice). Mice with two WT alleles and Cre expression were defined as wild-type mice (WT mice). Genotyping by tail preparation and PCR were performed at 2 weeks of age and the following experiments were carried out when the mice were 8 to 12-weeks old unless otherwise noted.

UUO-induced mouse model: Eight-week-old male mice were randomly divided into sham operation groups and unilateral ureteral obstruction (UUO) groups. Sham- operated mice were used as controls (n=15 per group). For the UUO model, sevoflurane was used to maintain general anesthesia for the convenience of skin preparation and surgical operation. After the surgical area was strictly disinfected, mice underwent ligation of the left ureter and were sacrificed 2 weeks later.

FA-treated mouse model: Eight-week-old male mice were randomly divided into a control group and an FA-treated group. For the FA model, mice were induced with a single intraperitoneal injection of FA (250 mg/kg, dissolved in 300 mM NaHCO3) and sacrificed 1 month later.

SB431542 treatment mouse model: Eight-week-old male mice were randomly divided into sham groups, UUO groups, and UUO+SB431542 groups. The sham group and UUO group were consistent with the processing described above. For the UUO plus SB431542 group, mice were treated with continual intraperitoneal injections of SB431542 (2 mg/kg/day, dissolved in 10 % PEG400 solution) for 10 days after UUO surgery and were sacrificed 4 days later.

Erlotinib treatment mouse model: Eight-week-old male mice were randomly divided into sham groups, UUO groups, and UUO+Erlotinib groups. The sham and UUO groups were consistent with the processing described above. For the UUO plus Erlotinib group, mice were administered with Erlotinib (50 mg/kg/day, suspended in sterile PBS before administration) once daily for 14 consecutive days via the oral route after the operation.

### Cell culture and treatments

HK-2 cells (Cell Bank, Chinese Academy of Sciences) were maintained in DMEM/F12 containing 10% FBS, penicillin (100 units/ml), streptomycin (100 μg/ml), and 1% L-glutamine unless otherwise noted. To detect the mRNA and protein level of *Glce*, HK- 2 cells (2 × 10^5^/well) were seeded in a 60 mm dish and then stimulated with 10 ng/ml TGF-β at different times. In the later experiment, HK-2 cells transfected with various plasmids were stimulated with or without TGF-β (10 ng/ml) for 24 h. To determine the action target, HK-2 cells were treated with TGF-β (10 ng/ml) for 24 h in the presence of SIS3 (5 μM) or SB431542 (10 μM).

### Statistical analysis

Statistical analyses for the study were performed using GraphPad Prism 8.0 software. All values are expressed as the standard error of the mean (SEM). 2-tailed Student’s t test was used to test for significance between the experimental and control groups. When 3 or more groups were assessed, 1-way ANOVA with Tukey’s or Dunnett’s multiple-comparison *post hoc* test were used. Correlation analyses were performed using Spearman’s correlation coefficient. *P* values < 0.05 are considered statistically significant.

Additional details for all methods are provided in the Supplementary Methods.

## DATA AVAILABILITY

All data are available in the main text or the supplementary materials.

## ACKNOWLEDGMENTS

We thank Shanghai Changzheng Hospital for providing us human renal biopsies.

## Author contributions

Xiaoqi Jing, Jun Wu and Jingru Ning contributed equally to the work and should be considered as co–first authors. The study was designed by Xiaoqi Jing, Kan Ding, Jun Wu and Changlin Mei; Experimental analysis was performed by Xiaoqi Jing and Jun Wu; Animal experiments were performed by Xiaoqi Jing and Zhenyun Du, which was assisted by Lulin Huang; Human renal immunohistochemical stains and related independent pathological evaluation were performed by Jun Wu and Ran Wang; Protein-protein docking analysis was performed by Xiaoyu Ding; Surface plasmon resonance assay was performed by Xiaoqi Jing and Xiaojiang Wang; Xiaoqi Jing and Jingru Ning wrote the manuscript; Kan Ding and Changlin Mei performed the critical revision and editing of the manuscript; All authors approved the final version of the manuscript.

## Funding

Shanghai Municipal Science and Technology Major Project, National Key R&D Program of China (2022YFA1303802)

National Natural Science Foundation of China (32271332, 31870801) High-level New R&D Institute (2019B090904008)

High-level Innovative Research Institute (2021B0909050003) from Department of Science and Technology of Guangdong Province Zhongshan Municipal Bureau of Science and Technology

---

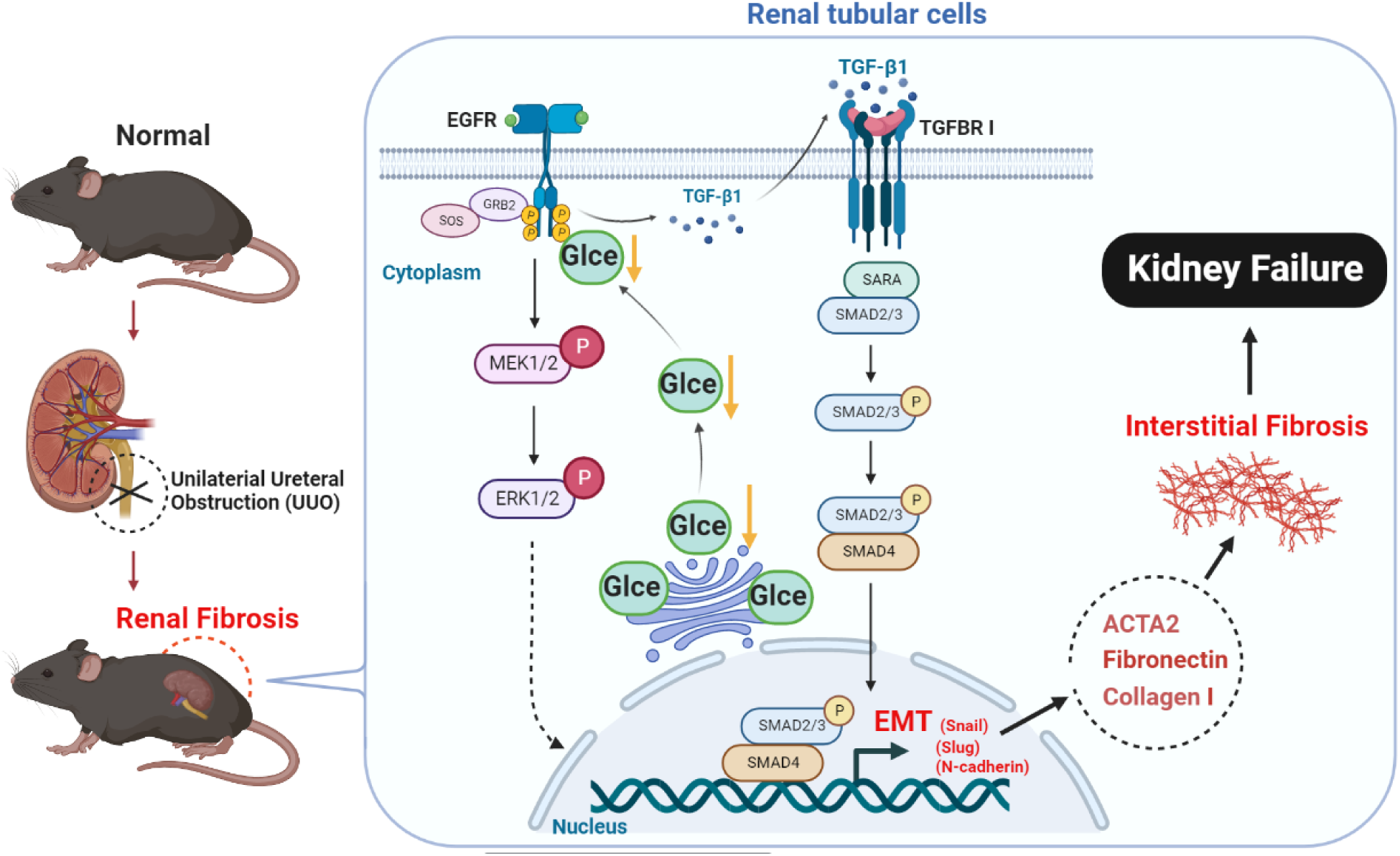

